# G-quadruplex DNA structures mediate non-autonomous instruction of breast tumour microenvironments

**DOI:** 10.1101/2023.01.16.524243

**Authors:** Pascal Hunold, Michaela N Hoehne, Martha Kiljan, Olivia van Ray, Jan Herter, Grit S Herter-Sprie, Robert Hänsel-Hertsch

## Abstract

Breast cancer is characterised by genetic and epigenetic alterations, such as G-quadruplex (G4) DNA secondary structures. Here, we uncover differentially enriched G4 structure-forming regions (∆G4Rs) and interlinked transcriptomes in the tumour microenvironment (TME) of breast cancer PDX models *in vivo*. We show that well-defined breast cancer cell models non-autonomously instruct ∆G4Rs and transcriptomes in the epigenomes of primary macrophages *in vitro*. Mechanistically, we uncover that TNBC secretes, amongst other factors, glucocorticoids to promote G4-linked activation of *octamer-binding transcription factor 1* (OCT-1) and thereby reprogramme macrophages into an immunosuppressed and immunosuppressive state. This epigenetic mechanism is of clinical importance since instructed macrophages selectively associate with the triple-negative breast cancer (TNBC) basal-like 2 (BL2) subtype and with the distinct TNBC molecular signature derived from 2,000 primary breast cancer samples. Altogether, our data suggest that G4 formation is not only prevalent in breast cancer genomes but relevant in their TMEs as well, which is of clinical importance for cancer stratification and the discovery of novel actionable drivers.

## Introduction

DNA stretches of adjacent guanines can self-assemble into four-stranded DNA secondary structures called G-quadruplexes (G4s)^1,2^. We revealed prevalent G4 formation in nucleosome-depleted regions (NDRs) of highly expressed genes in mutation hotspots of cancer genomes^3,4,5^. Besides cancer, human and murine pluripotent states have displayed higher G4 levels in regulatory regions relative to differentiated states^6,7^. However, the physiological role of G4s under healthy conditions remains elusive in mammals. Transcription factors (TFs) can establish and change the location of NDRs. Numerous TFs demonstrated similar binding preferences to endogenous G4s with respect to their consensus binding motifs^8,9^. Hence, many TFs may interact with G4s to regulate transcription. Previously, we profiled G4 landscapes of tumours from 22 breast cancer patient-derived xenograft (PDX) models. We found a remarkable association with breast cancer-derived TF programmes, suggesting that G4s are valuable proxies to infer TF activity in clinically-relevant cancer samples^10^. Clinically, breast cancer has been divided into four subtypes according to oestrogen and progesterone receptor (collectively called hormone receptors (HR)) as well as human epidermal growth factor receptor 2 (HER2) status. Triple-negative breast cancer (TNBC) represents the subtype of the poorest clinical prognosis, displaying at least four clinically relevant subtypes (BL1, BL2, M and LAR) ^11,12,13,14^. Integrating somatic copy number alteration with transcriptomic data of primary tumours from 2,000 breast cancer patients has established at least ten distinct molecular subtypes, here further referred to as integrative clusters (ICs)^15^. Integration of quantitative G4-ChIP-seq data of tumours from breast cancer PDX models with gene signatures distinct to ICs revealed high intra-tumour heterogeneity for most PDX models. Importantly, quantitatively assessed G4 levels of PDX tumours correlated with their susceptibility to pharmacological G4 stabilising compounds^10^. The tumour microenvironment (TME) is progressively recognised as a critical aspect of disease progression and therapy responsiveness^16^. Large-scale single-cell and spatial omics analysis of TMEs of primary breast cancers revealed their association with genomic features and clinical outcomes^17^. The cellular portion of the TME is mainly composed of stromal cells, blood vessel linings, and infiltrating immune cells^18^.

Here, we uncover ∆G4Rs and altered transcriptomes in the TME of breast cancer PDX models *in vivo*. We show that well-defined breast cancer cell models non-autonomously induce ∆G4Rs and thus alter transcriptomes in primary macrophages *in vitro*. Mechanistically, we uncover that TNBC secretes, amongst other factors, glucocorticoids to promote G4-linked activation of OCT-1 and thereby reprogrammes macrophages into an immunosuppressed (major histocompatibility complex (MHC) II low) and immunosuppressive (*arginase 1* high) state. This epigenetic mechanism is of clinical importance since instructed macrophages selectively associate with the TNBC-BL2 subtype and with the distinct TNBC molecular signature derived from 2,000 primary breast cancers. Altogether, our data suggest that G4 formation is not only prevalent in breast cancer genomes but also relevant as an epigenetic feature in the TME, which is of clinical relevance for cancer stratification and discovering novel actionable drivers of the TME.

## Results

### Breast cancer PDX models establish distinct G4 landscapes and transcriptomes within their microenvironments

Upon transplantation of human breast tumours into genetically identical murine avatars (PDX), cells of the human TME are replaced with murine cells^19^. Previously, we found ΔG4Rs in the human cancer cell fraction across a panel of 22 breast cancer PDX models by qG4-ChIP-seq^10^. To reveal ΔG4Rs in the TME of the PDX qG4-ChIP-seq data, we analysed the PDX qG4-ChIP-seq data and selected reads that mapped to the murine genome (Fig. 1a). Importantly, we noticed that up to ∼20% of the PDX reads aligned to the murine genome, suggesting that murine TMEs adopt G4 DNA structures *in vivo*. Indeed, we revealed the existence of ΔG4Rs in 9 of 22 of the PDX-derived TMEs (Fig. 1b, Supplementary Fig. 1a). Independent of the TME sequencing fraction, we unravelled ∼120-800 ΔG4Rs for the nine PDX models (Fig. 1b, c, Supplementary Fig. 1a). This up to ∼6-fold difference in ΔG4Rs across the TMEs suggests that they adopt distinct epigenomic states, depending on the diverse tumour intra-heterogeneity of the PDX models. Of note, all PDX models were propagated in genetically identical mouse avatars of the same sex and age to prevent avatar-related effects on TME and cancer cells^10,19^. Pairwise comparison with hierarchical clustering of the ΔG4Rs revealed distinct TME subgroups regardless of the HR or IC status of the respective PDX models (Fig. 1d). In line with existing reports that endogenous G4 regions are prevalent in promotors of actively transcribed genes^3^, we revealed a striking fraction of TME ΔG4Rs at promoters (upstream and 5’UTR) of highly expressed genes (Fig. 1e, Supplementary Fig. 1b). Hierarchical clustering of differentially expressed genes revealed differences in the expression of myeloid cell-associated genes (*Fth1, Mcl1, Ly6e, Tgfbi*, and *Selplg*), genes associated to endocrine tissues (*Dchs1, Rgs5*, and *Lrrc58*) as well as stromal tissue genes (*Nkd2, Moxd1, Scn7a*, and *Col8a1/2*) (Fig. 1f, Supplementary Fig. 1c)^20^. Moreover, we observed substantial differences in the expression of genes associated with angiogenesis (*Sptbn1*), DNA damage repair and replication (*Herc2*), and autophagy (*Trp53inp2*)^20,21,22^ (Fig. 1f). We then hypothesised that the differences in G4 and transcriptional landscapes of our TMEs might arise from the underlying intra-tumour heterogeneity of PDX models. Here, we propose that different breast cancer subtypes underlying PDX models either attracted distinct TME-related cell types or instructed the same cells of the TMEs to acquire different epigenomes. To obtain broader insights into their respective cell compositions, we analysed the TME-related RNA-seq data of the PDX models. To assess these differences, we employed Consensus^TME^, a state-of-the-art integrative bioinformatics tool, to determine the relative abundance of immune cells from gene expression data^23^. In line with the immunosuppressed phenotype of NSG mice, we found a depletion of lymphocytes across the PDX models (Fig. 1g)^24^. Importantly, we observed a consistent enrichment of stroma and myeloid cells among the PDX models (Fig. 1g). However, we observed no substantial differences in the composition of TME-related cell types across the PDX models (Fig. 1g). Our data suggest that cancer cells of the different PDX models instruct distinct epigenetic alterations into the same prevalent TME-related cell types, such as macrophages.

**Fig. 1.**
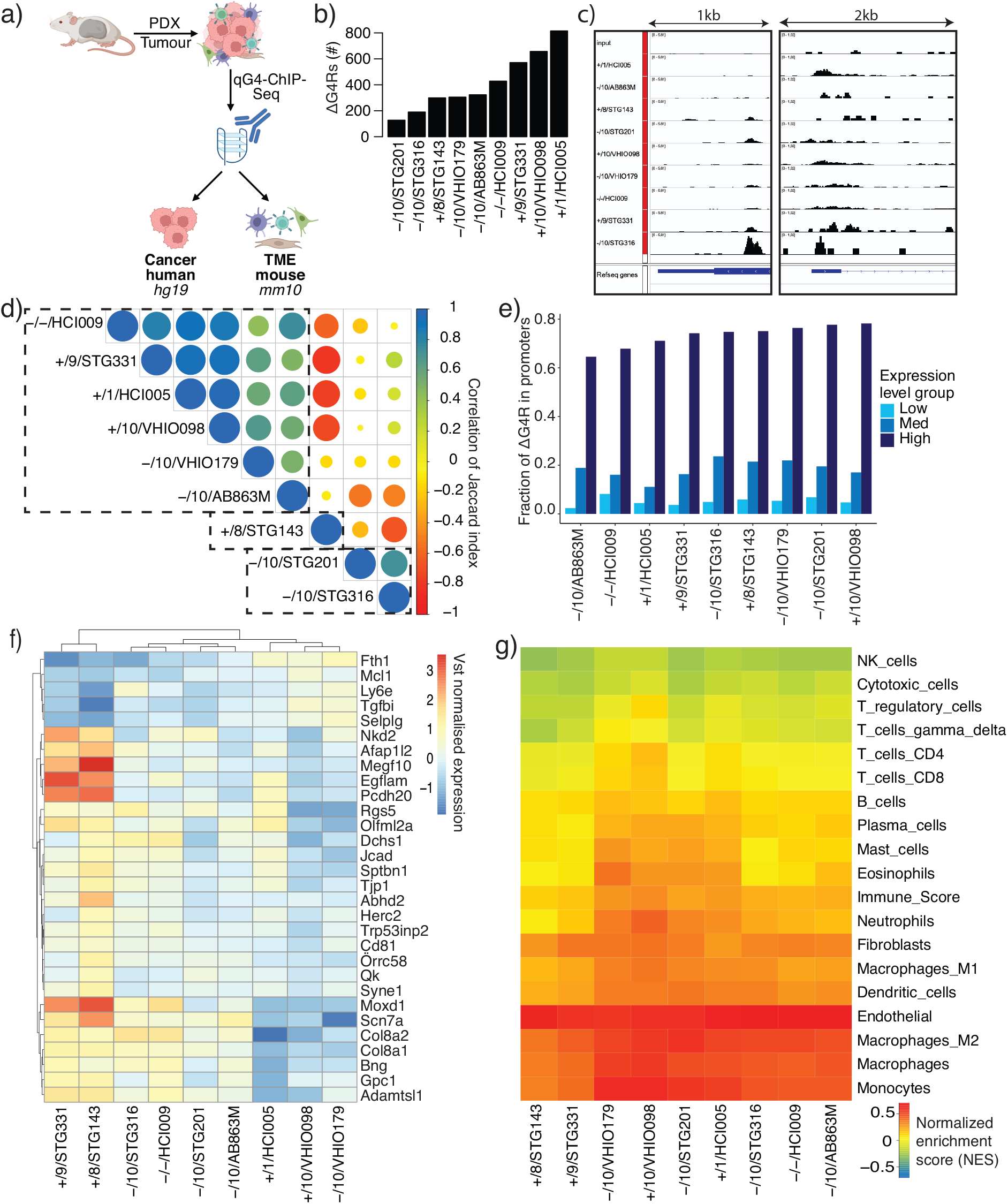
Breast cancer PDX models establish distinct G4 landscapes and transcriptomes within their microenvironments. **a** Scheme of quantitative G4-ChIP-seq experiment and origin of human and murine sequencing reads. TME: tumour microenvironment, hg19 (*H. sapiens* genome), mm10 (*M. musculus* genome). **b** Number of mm10 ΔG4Rs for each PDX model. **c** Genome browser views (IGV) of MN ΔG4Rs. **d** Heatmap visualising the similarity of ΔG4Rs across PDX models. **e** The x-axis shows high (blue), medium (red) and low (black) levels of gene expression. The y-axis shows the percentage of ΔG4Rs in the expressed promoters for each PDX. **f** Heatmap of the 30 most variable genes for all 9 PDX models. Mean vst value shown for each PDX model. **g** ConsensusTME heatmap. The heatmap shows the normalised enrichment score for each TME cell type of each PDX model. From RNA-seq data, mean counts per million were calculated for each PDX model and used as input for Consensus^TME^.

### Non-autonomous instruction of ΔG4R-related TF activity in macrophages by different breast cancer subtypes

We recently showed that ΔG4Rs of breast cancer PDX models revealed novel TF programs^10^. To explore whether different breast cancer subtypes can instruct distinct epigenetic mechanisms in our TMEs (Fig. 1g), we studied ΔG4Rs and related TF activities in primary murine bone marrow-derived macrophages (BMDMs) *in vitro* (Fig. 2a). We incubated BMDMs with the medium conditioned by well-characterised breast cancer cell lines, representing either HR^+^ (MCF-7) or TNBC (MDA-MB-231) (Fig. 2a). Strikingly, only the medium conditioned by the TNBC cell line restored the proliferative capacity of terminally differentiated BMDMs (Fig. 2a). Comparative analysis of G4 landscapes in treated BMDMs identified ∼5.000 HR^+^ and TNBC-specific ΔG4Rs, demonstrating that different breast cancer subtypes non-autonomously instruct distinct epigenetic mechanisms in macrophages (Fig. 2b, c, Supplementary Fig. 2a). Our findings suggest that only the TNBC but not the HR^+^-defined cell line released soluble factors into their medium to instruct ΔG4Rs into terminally differentiated macrophages to provoke a novel proliferation phenotype. To investigate whether the TNBC-instructed G4 landscape in macrophages originated from the activity of particular regulatory programmes, we used TF occupancy prediction by analysing mapped G4 instead of chromatin accessibility signals (Fig. 2d, Supplementary Fig. 2b)^27^. ΔG4R-based TF occupancy of TNBC-treated relative to untreated macrophages uncovered factors involved in the canonical and non-canonical NFκB pathway (NFKB1, NFKB2, REL, RELA, and RELB) and myeloid cell differentiation-associated TFs (CEBPA, CEBPB, CEBPE, and CEBPG). We validated predicted changes in ΔG4R-related TF activity via RT-qPCR and RNA-seq. For example, we showed notable induction (>6-fold change) of the NFκB target genes *Tnfa* and *Cxcl1* in TNBC-treated BMDMs (Supplementary Fig. 2C). Intriguingly, several members of the *Pit-Oct-Unc 2* (*POU*)-family were exceptionally enriched in TNBC-treated BMDMs (Fig. 2d). Most *POU*-family members have been reported to be involved in diverse transcriptional programmes, with *POU5F1* (OCT-4) as the best-characterised representative^26,27^. Although *POU*-family members act in different epigenetic programmes, all of which are suggested to play crucial roles in cell identity-defining processes, such as directed differentiation or maintenance of stemness^27^. To identify which *POU*-family members are active in TNBC-treated macrophages, we examined the activation of their reported target genes (Supplementary Fig. 2d). We found target genes of the *POU2* subfamily (*POU2F1* and *POU2F3*) significantly altered. In contrast, targets of the *POU3* subfamily (*POU3F1* and *POU3F2*) and *POU5F1* remained unchanged. Intriguingly, OCT-1 (*POU2F1*) exhibited increased chromatin binding and nuclear localisation in TNBC-treated compared to untreated BMDMs (Fig. 2e, f). OCT-1 has been reported as a prognostic marker linked to poor clinical outcomes in various cancers^28,29,30,31,32^. However, the role of OCT-1 within the TME is not known. In support of a TNBC-instructed role of OCT-1 in macrophages, the OCT-1 target gene R-ras – a member of the Ras family – is significantly higher expressed (p-value ∼9.9^-20^) in our TNBC-treated relative to untreated BMDMs^33,34^. R-ras has been reported to enhance cell cycle progression and DNA synthesis leading to increased proliferation^33^. Furthermore, R-ras is required to activate cell adhesion via αMβ2 integrin in macrophages^33,35^. *ITGAM* (αMβ2 integrin) is higher expressed in our TNBC-treated in comparison to untreated macrophages (p-value ∼1.2^-10^). Cyclin D1 is expressed in most tissues except lymphoid and myeloid cells^36^. In addition, the cell cycle regulator cyclin D1 (*Ccnd1*) – another OCT-1 target gene^37,38^ – is upregulated in our TNBC-treated relative to BMDMs (p-value ∼8.3^-15^). We further found that TNBC-instructed ΔG4R-related OCT-1 footprints from primary macrophages are notably more enriched in highly expressed genes of TMEs from the PDX models (Supplementary Fig. 2e). We hypothesised that TNBC instructs an immunosuppressed and immunosuppressive phenotype through the release of soluble factors that activate OCT-1 in macrophages. Glucocorticoid receptors can recruit OCT-1 to chromatin upon activation^39^. In support of our observations in primary macrophages, we revealed substantial activation of OCT-1 in the macrophage line RAW246.7 when treated with hydrocortisone (Fig. 2g, h). We observed non-autonomous instruction of differential G4 landscapes in macrophages by two distinct breast cancer subtypes. We revealed G4-related OCT-1 activation in TNBC-instructed macrophages and independently recapitulated this with hydrocortisone treatment using an established macrophage line. These findings suggest that TNBC releases glucocorticoids in the TME to reprogram macrophages into an immunosuppressed and immunosuppressive phenotype.

**Fig. 2.**
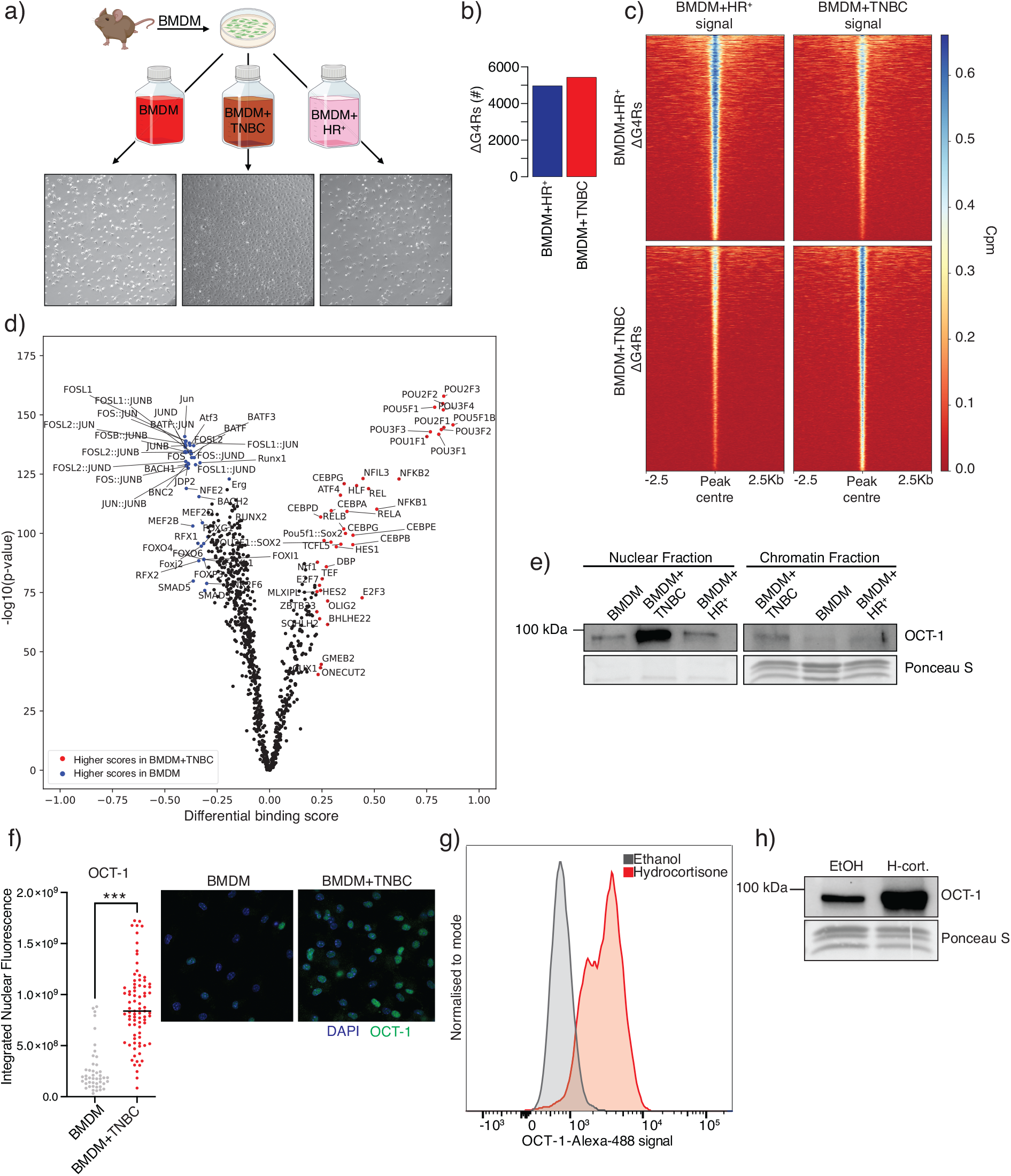
Non-autonomous instruction of ΔG4R-related TF activity in macrophages by different breast cancer subtypes. **a** Experimental scheme (Created with BioRender.com) with phase contrast microscopy images. Images were acquired with 10X magnification using an Olympus IX83. **b** Numbers of ΔG4Rs found in BMDMs treated with conditioned media either from HR^+^ (BMDM+HR^+^) or TNBC (BMDM+TNBC) cells. **c** Heatmap using ΔG4Rs of BMDM+TNBC and BMDM+HR^+^ and the signals (Cpm normalised bigwig) for both. **d** TOBIAS results. The Y-axis of scatterplots shows the significance, and the x-axis shows the differential binding score. Footprints in reads (bam files) were analysed in ΔG4Rs of BMDM or treated with conditioned media from TNBC (BMDM+TNBC). **e** Western blot of soluble and chromatin-bound nuclear fractions of BMDM, BMDM+HR^+^ and BMDM+TNBC conditions. OCT-1 signal and Ponceau (loading control). **f** Immunofluorescence of OCT-1 in BMDM and BMDM+TNBC conditions. Left: quantification of integrated nuclear fluorescence. *** p-value < 0.001. Right: representative images. **g** RAW246.7 were treated with 15 ng/mL hydrocortisone (or ethanol as solvent control) for 24 h. Representative histogram plots indicate nuclear OCT-1 protein (n=2). **h** Western blot of chromatin-bound OCT-1 in RAW246.7 treated as in **g**. Ponceau S serves as a loading control.

### TNBC-instructed macrophages exhibit an immunosuppressed and immunosuppressive phenotype of clinical relevance

To comprehensively understand how reprogramming of the BMDM epigenome via distinct signalling pathways and TF families could benefit TNBC, we examined the gene ontologies of TBNC-treated and untreated BMDMs (Supplementary Fig. 3a). In line with the immunological defence function of macrophages, diverse immune processes and responses to noxious signals were the most altered ontologies. Strikingly, TNBC-instructed BMDMs showed a strong down-regulation of mechanisms associated with MHC II presentation and processing. In support of this, *Cd300e*, a member of the Cd300 family, is significantly higher expressed (p-value ∼4.7^-18^) in our TNBC-treated relative to untreated macrophages^40^. In line with the literature^41^, the increased expression of Cd300e leads to the downregulation of *CIIta* (p-value ∼9^-85^) and, thereby, loss of MHC II expression, such as *Cd74* (p-value ∼1.6^-52^), *H2-Aa* (p-value ∼2.2^-144^), and *H2-Ab1* (p-value ∼1.3^-60^) in our TNBC-treated BMDMs (Fig. 3a, Supplementary Fig. 3b). To validate the immunosuppressed phenotype of TNBC-treated BMDMs, we analysed the levels of MHC II proteins in TNBC-treated vs untreated BMDMs by immunofluorescence microscopy. We observed decreased MHC II level in TNBC-treated BMDMs (Fig. 3b). Moreover, the M2 macrophage marker arginase 1 *(Arg1)* is substantially increased in our TNBC-treated BMDMs (p-value ∼1.2^-18^) (Fig. 3c)^42^. Importantly, *Arg1* was significantly (FDR ∼4.4^-6^) higher expressed in the TME of the PDX models that stratify to IC10 compared to non-IC10 models (Fig. 3d), supporting the possibility that our TNBC-related observations *in vitro* are of clinical relevance for TNBC. To strengthen the clinical implication of the macrophage immunosuppressive and immunosuppressed phenotype, we associated the gene expression differences of TNBC-treated BMDMs with the gene signatures distinct for each of the ten integrative clusters (ICs) of breast cancer^10^. Of note, only differentially expressed genes of TNBC-treated BMDMs were consistently associated with the IC10 gene signature, the only IC related to primary TNBC, using two independent association strategies (Fig. 3e, Supplementary Fig. 3c, d). Multiomics-based stratification of 192 breast cancer patients confirmed the existence of four TNBC subtypes, *i*.*e*., BL1, BL2, LAR and M^13^. Strikingly, we found a distinct enrichment of BL2 patients with the top 100 differentially upregulated genes (NES ∼2.2, p-value < 0.0001) of TNBC-treated BMDMs (Fig. 3f). Importantly, we noticed that the transcriptomes of BL2 patients are also enriched for the G4-linked OCT-1 target genes of our TNBC-treated BMDMs (Fig. 3f). Therefore, we anticipated that a considerable amount of BL2 patient tumours were infiltrated with immunosuppressed macrophages (Fig. 3g). Indeed, we found a substantial amount of BL2 patients (cluster 1, 11/37) that exhibited depletion of MHC-II related gene expressions (Fig. 3g). In summary, our findings suggest that TNBC can actively instruct the epigenome of tumour-associated macrophages to promote a G4-OCT-1-linked proliferative, immunosuppressive and immunosuppressed phenotype in TNBC patients of the BL2-subtype.

**Fig. 3.**
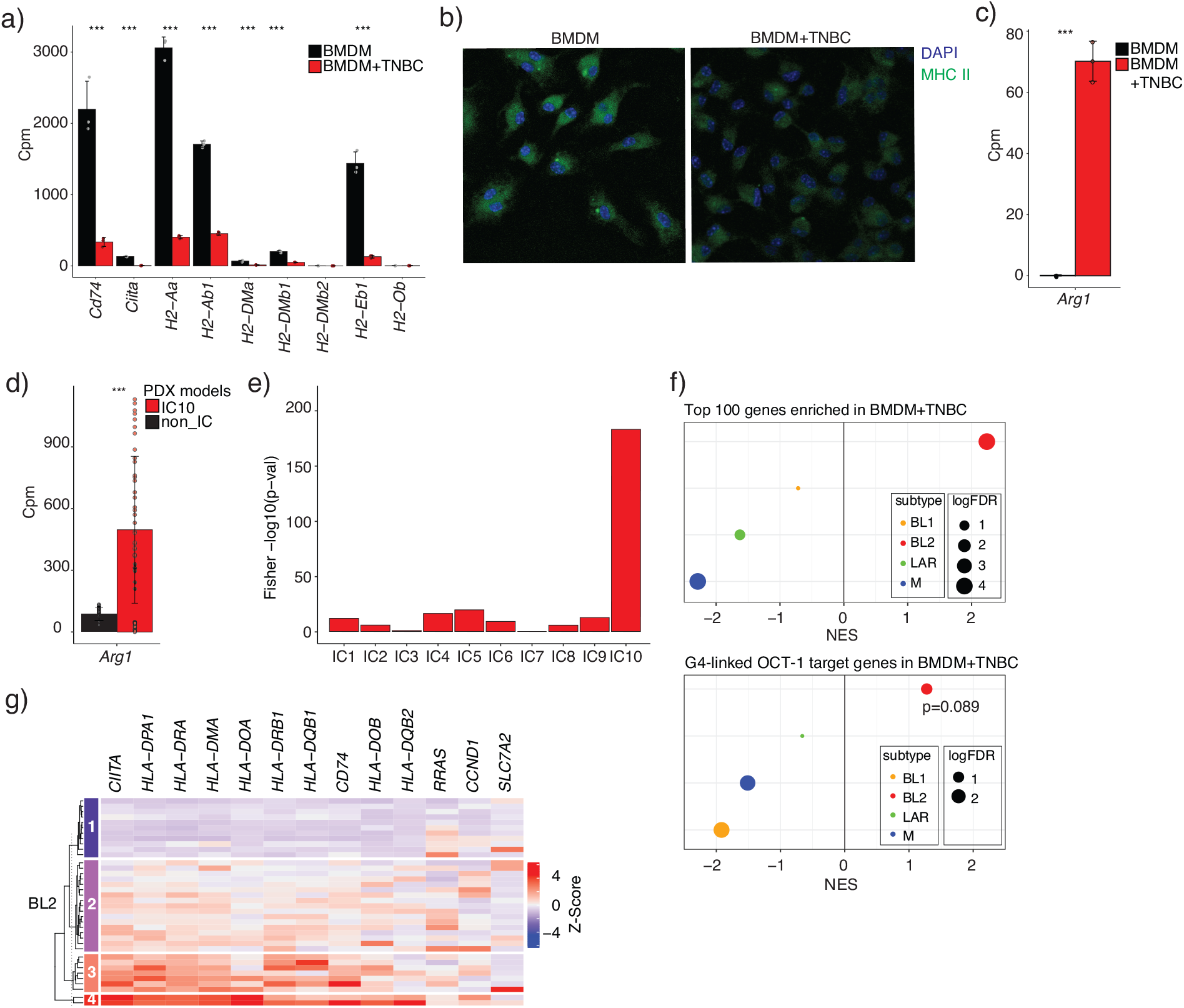
TNBC-instructed macrophages exhibit an immunosuppressed and immunosuppressive phenotype of clinical relevance. **a** RNA-seq mean cpm values and standard deviation are shown for MHC II subunits in BMDM and BMDM+TNBC conditions. *** p-value < 0.001, from the differential analysis. **b** Immunofluorescence of MHC II and DAPI in BMDM and BDMD+TNBC conditions. **c** RNA-seq mean cpm values and standard deviation are shown for *Arg1* in BMDM and BMDM+TNBC conditions. *** p-value < 0.001, from the differential analysis. **d** RNA-seq mean cpm values and standard deviation are shown for *Arg1* in PDX models stratified by IC10 status. *** p-value < 0.001 from the differential analysis. **e** -log10(p-value) of Fisher’s exact test extracted from overlaps of the integrative cluster (IC) signature genes (molecular subtypes of breast cancer) with differentially enriched genes revealed in BMDM+TNBC. **f** GSEA preranked of TNBC subtypes and top 100 enriched genes in BMDM+TNBC vs BMDM or OCT-1 targets identified in BMDM+TNBC using TOBIAS as gene set, respectively. **g** Heatmap of BL2 subtype patients showing genes identified in this study, Kmeans=4.

## Discussion

Until now, prevalent G4 DNA secondary formation has primarily been described as a cancer phenomenon to promote genome instability and altered gene regulation^3,8,9,10, 43,44^. Now and for the first time, we reported differentially enriched G4 formation within the microenvironment of breast cancer PDX models *in vivo*^10,19^. Transcriptome-based cell identification analyses showed no substantial changes in the cell composition of TMEs across the different PDX models (Fig. 1g). Importantly, our findings suggest that different breast cancer subtypes can instruct distinct G4 landscapes and related epigenetic alterations in the same TME-related cell type. Here, we revealed non-autonomous instruction of distinctive G4 landscapes in primary murine macrophages by two most different well-characterised human breast cancer cell models, reflecting TNBC or HR^+^. We discovered that the TNBC but not the HR^+^ model instructed an extraordinary proliferation phenotype in such macrophages. This TNBC-instructed phenotype was accompanied by distinct G4 landscape alteration related to a substantial increase in OCT-1 activity in primary macrophages as well as the TME of the PDX models (Fig. 2d, Supplementary Fig. 2e). Transcriptomic analyses displayed a considerable induction of the previously described OCT-1 targets R-ras and cyclin D1 to drive cell cycle progression and thereby cell growth^45^. We furthermore revealed that this G4-OCT-1-linked proliferation phenotype is heavily compromised in its MHC II processing and presentation. In support of this immunosuppressed macrophage phenotype, our gene expression analysis revealed TNBC-instructed abrogation of the MHC II master regulator CIIta (Fig. 3a). Intriguingly and in line with this MHC II repression, OCT-1, CEBP, RELA (activated in TNBC-treated BMDMs) all share proximal response elements with nuclear receptors like the glucocorticoid receptor (GR)^46^. GR has been reported to alter the expression of *IL-10, Ccl5, IL-1β, IL-6, IL-12a*, and *Tnfa*, all showing significantly changed expression in TNBC-instructed macrophages alongside *GR*^47^. Glucocorticoids can screw macrophages towards M2 polarisation^48^. In line, TNBC-instructed macrophages showed drastically increase in the expression of M2 marker gene *arginase 1*, which is associated with poor clinical prognosis. Arginase 1 is known to deplete arginine locally from the TME to suppress further other immune cells like T cells^49,50,51^. High expression of GR in breast cancer is linked to the development of TNBC and poor clinical outcome^52,53^. The pharmacological antagonism of GR-sensitised TNBC against DNA-damaging chemotherapy was indicated as a new therapeutic approach^54^. We have shown that the treatment of macrophages with hydrocortisone – a pharmacologically used glucocorticoid – provokes substantial recruitment of OCT-1 to the chromatin of macrophages (Fig. 2g, h). Importantly, we showed that this TNBC-induced G4-linked macrophage phenotype is of clinical importance since an independent TNBC-related gene signature (IC10) derived from the genomic and transcriptomic analysis of 2,000 primary breast cancers associated explicitly with our macrophage phenotype.

Moreover, we found a significant upregulation of arginase 1 in the TME of TNBC-IC10 PDX models compared to non-IC10 models. This transcriptome alteration further highlights the ability of TNBC to reprogramme its TME *in vivo*. A recent multiomics study of 192 TNBC patients confirmed the existence of at least four distinct subtypes^13^. We revealed a significant and selective association of our TNBC-instructed G4-linked macrophage phenotype in TNBC BL2 breast cancer patients. We propose the existence of a novel TNBC BL2 subtype that is highly infiltrated with immunosuppressed and immunosuppressive macrophages.

In conclusion, our work indicates a novel mechanism of TNBC to non-autonomously instruct a highly proliferative G4-linked OCT1 immunosuppressed and immunosuppressive phenotype in macrophages (Fig. 4). We provided evidence that non-autonomous instruction of TNBC may originate from the secretion of – amongst others - glucocorticoids into its TME. We propose the therapeutic potential to employ GR antagonism against TNBC in G4-OCT1-linked BL2-subtyped patients to re-establish an immunogenic TME by reversal of cancer-instructed epigenetics in residential macrophages.

**Fig. 4.**
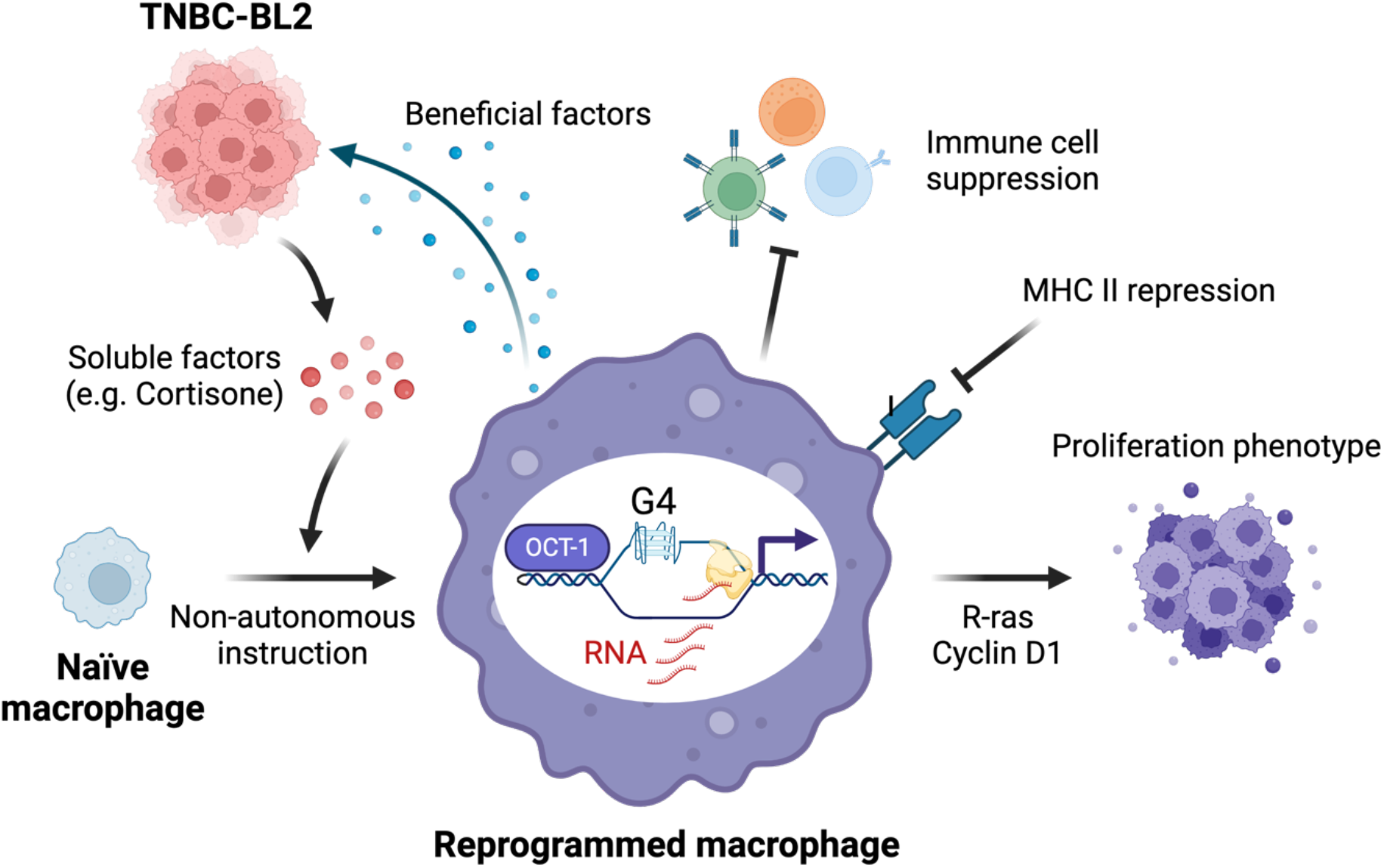
TNBC-BL2 reprogrammes macrophages via G4-OCT-1 activation. A model (created with BioRender.com) displaying epigenetic reprogramming of naïve macrophages by TNBC-BL2. Upon release of soluble factors (e.g., cortisone) by the TNBC-BL2, naïve macrophages adopt G4 DNA secondary structures in regions where OCT-1 promotes an immunosuppressed (low *MHC II*), immunosuppressive (high *Arg1*) and highly proliferative (*R-ras, Ccnd1*) gene expression programme. As cells with high secretory activity, reprogrammed macrophages could release cancer-benefitting factors to support TNBC-BL2 growth.

## Material and methods TME

### Cell culture

MCF-7 were kindly provided by Katrin Paeschke (Department of Oncology, Hematology and Rheumatology, Bonn, Germany). MDA-MB-231 were kindly supplied by Thorsten Stiewe (Institute of Molecular Oncology, Marburg, Germany). RAW246.7 were kindly provided by Stephan Baldus (Center for Molecular Medicine Cologne, Cologne, Germany). All cell lines were cultured in Dulbecco’s Modified Eagle Medium (Gibco, DMEM, cat. no. 31966-021) supplemented with 10 % FBS (Gibco) at 37 °C in 5 % CO_2_. All cell lines were tested negative for mycoplasma MycoSPY Mycoplasma PCR Detection Kit (Biontex, cat. no. M030-050).

### Murine BMDM isolation and differentiation

Mice were bred and maintained in specific pathogen-free conditions in the animal facility Weyertal at the University Hospital Cologne. The local animal care committee approved all animal experiments. C57/BL6J mice were sacrificed between 8 and 16 weeks by cervical dislocation, and both the femur and tibia were dissected. After the remaining tissue was dehydrated with 70% ethanol, bones were flushed with DMEM + 10% FBS + 1% P/S (Penicillin/Streptomycin) with a 10 mL syringe with a 23-gauge needle. Erythrocytes were lysed with 1 ml ACK buffer for five minutes at room temperature. The cells were washed, and 5×10^6^ cells were plated in 5 ml DMEM + 10% FBS + 1% P/S + 20 ng/ml recombinant mouse Macrophage Colony-Stimulating Factor (Biolegend, cat. no. 576402) onto 60 mm tissue culture dishes. Cells were cultured at 37 °C in 5 % CO_2_. Macrophages were ready for downstream applications after seven days.

### Cancer-conditioned media treatment

DMEM (Gibco, DMEM, cat. no. 31966-021) supplemented with 10 % FBS (Gibco) were conditioned by 1×10^7^ plated MCF-7 or MDA-MB-231 cells for 24 h. Cancer cells and debris were removed by sterile filtering of the conditioned media. BMDMs were incubated for 24 h with MCF-7 (HR^+^) or MDA-MB-231 (TNBC) conditioned medium, respectively. Untreated BMDMs were incubated with fresh DMEM.

### RNA isolation and RNA-seq

Total RNA from BMDMs were extracted using the NucleoSpin RNA Kit (Macherey-Nagel, cat. no. 740955.50) according to the manufacturer’s protocol. RNA-seq libraries were generated with the QuantSeq 3’ mRNA-Seq Library Prep Kit FWD with Unique Dual Indices for Illumina (Lexogen, cat. no. 115.384) and sequenced on an Illumina NovaSeq 6000 platform using NovaSeq 6000 SP Reagent Kit v1.5 (100 cycles) (Illumina, cat. no. 20028401). All RNA-seq experiments were conducted in three independent technical replicates.

### Reverse Transcription and RT-qPCR

Isolated total RNA (see above) was reverse transcribed using the High-Capacity cDNA Reverse Transcription Kit (Applied Biosystems, cat. no. 4368814) according to the manufacturer’s protocol. RT-qPCR was performed on a Bio-Rad CFx96 Touch Real-Time PCR Detection System using Fast SYBR Green Master Mix (Applied Biosystems, cat. no. 4385612). Ct values were analysed using the ΔΔCt method with *Tbp* as housekeeping genes.

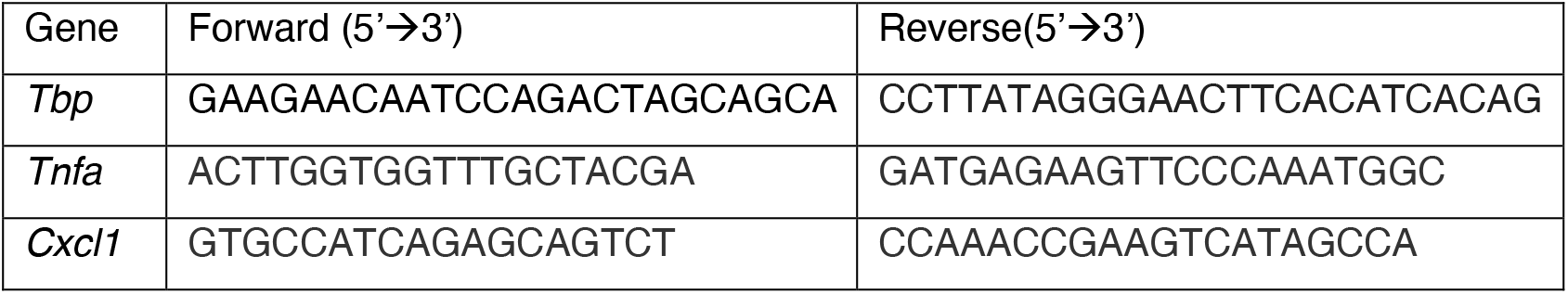

### Cleavage Under Targets and Tagmentation (CUT&Tag)

CUT&Tag experiments were performed as previously described^55^ with the following modifications. Instead of immobilisation of nuclei on concanavalin A-conjugated beads, nuclei were washed by centrifugation at 600 xg for 3 min at RT. For G4-CUT&Tag, 1×10^5^ fixed nuclei were incubated with 60 nM anti-G4 BG4 antibody^56^ on a nutating platform at 4 °C overnight. The next day, nuclei were first incubated with 0.5 µg anti-FLAG antibody (Cell Signaling, cat. no. 2368) for 30 minutes at room temperature, followed by an additional 30 min incubation at room temperature with 0.5 µg anti-rabbit IgG antibody (NOVUS BIOLOGICALS, cat. no. NBP1-72763). Libraries were paired-end sequenced on an Illumina NovaSeq 6000 platform using NovaSeq 6000 SP Reagent Kit v1.5 (100 cycles) (Illumina, cat. no. 20028401) with 0.5 % PhiX Control V3 (Illumina, cat. no. FC-110-3001). All CUT&Tag experiments were performed in three independent technical replicates.

### Protein A-Tn5 purification and transposome preparation

To use the formerly described protein A-Tn5 for CUT&Tag^55^ in combination with the FLAG-tagged anti-G4 BG4 antibody, the triple FLAG tag was removed from the 3Xflag-pA-Tn5-Fl plasmid (Addgene, #124601) by molecular cloning. For molecular cloning, the 3Xflag-pA-Tn5-Fl plasmid was amplified using mutagenesis primers 5’-ACCATGGGTATGACCATGATTACGCC-3’ and 5’-CATGGTCATACCCATGGTATATCTCCTTC-3’ and cloned with the In-Fusion HD Cloning Plus (Takara Bio, cat. no. 638911). Protein A-Tn5 was expressed and purified as previously described^55^. For transposome preparation, 16 µL of 100 µM annealed Tn5MEDS-A and Tn5MEDS-B oligonucleotides were mixed with 100 µL of proteinA-Tn5 glycerol stock and incubated for 50 min at 23 °C ^55^. The prepared transposome was stored at -20 °C.

### BG4 antibody expression and purification

The anti-G4 BG4 antibody was expressed and purified as formerly described^43^. In short, with pSANG10-3F-BG4 (Addgene, #55756), transformed *E. coli* Rosetta2 (DE3) were cultured in 2xTY medium supplemented with 50 µg/mL kanamycin (Alfa Aesar, cat. no. J17924). To extract periplasmatic BG4, bacteria were incubated with TES buffer on ice for 15 min. Upon pelleting bacteria by centrifugation at 8000 xg for 10 min at 4 °C, extracted BG4 antibody was bound to HIS-Select Nickel Affinity Gel (Sigma Aldrich, cat. no. P6611) for 1 h on a rolling shaker at RT. Affinity purified BG4 antibody was eluted with 250 mM imidazole (Sigma Aldrich, cat. no. I2399) and dialysed for 48 h at 4 °C against PBS. The purity of the BG4 antibody was assessed by SDS-PAGE and Coomassie blue staining. Protein concentration was determined with Qubit Protein Assay Kit (Invitrogen, cat. no. Q33212), adjusted to 3 µM with PBS and stored at -80 °C.

### Western Blot of subcellular fractions

Western blot analysis of subcellular fractions was performed as previously described^57^. Briefly, cells were first lysed in E1 buffer (50 mM HEPES-KOH pH 7.5, 140 mM NaCl, 1 mM EDTA, 10 % glycerol, 0.5 % NP-40 alternative, 0.25 % Triton X-100, 1 mM DTT, 1X cOmplete Proteinase Inhibitor) to isolate the cytoplasmatic fraction. Subsequently, soluble nuclear proteins were extracted in E2 buffer (10 mM Tris pH 8.0, 200 mM NaCl, 1 mM EDTA, 0.5 mM EGTA, 1X cOmplete Proteinase Inhibitor). Chromatin-bound proteins were isolated by incubation with ∼500 U Benzonase (Sigma Aldrich, cat. no. E1014) for 20 min at room temperature in E3 buffer (500 mM Tris pH 6.8, 500 mM NaCl, 1X cOmplete Proteinase Inhibitor). To increase the isolation efficiency of chromatin-bound proteins, chromatin was sonicated for five cycles (30 sec ON, 30 sec OFF) on a high setting of a Diagenode Bioruptor plus before Benzonase digest. Debris from all fractions was removed by centrifugation for 10 min at 16,000 xg and 4 °C. Protein concentration was assessed with the Qubit Protein Assay Kit (Invitrogen, cat. no. Q33211). 20 µg protein were separated in Mini-PROTEAN TGX Precast Protein Gels (Bio-Rad, cat. no. 4561086) and transferred onto nitrocellulose membranes (Bio-Rad, cat. no. 1704270). Total proteins were visualised by Ponceau S (Carl Roth, cat. no. 5938.1) and served as a loading control. After blocking the membrane in 5 % milk in TBS-T (20 mM Tris pH 7.0, 150 mM NaCl, 0.1 % Tween 20), membranes were incubated 1:1000 diluted anti-OCT1 (Invitrogen, cat. no. MA5-41265) overnight at 4 °C. The next day, membranes were washed three times with TBS-T and incubated with 1:2,500 diluted anti-rabbit-HRP secondary antibody (Invitrogen, cat. no. 32460) and visualised with Clarity Max Western ECL Substrate (Bio-Rad, cat. no. 1705062). All images were acquired on the Bio-Rad ChemiDoc MP Imaging System.

### Immunofluorescence and Image acquisition

Cells were fixed for 10 min in 4% formaldehyde (Carl Roth, cat. no. 4235.1) at room temperature. For intracellular staining against OCT-1, cells were permeabilised with 0.1% Triton X-100 in PBS for 15 min at room temperature. For cell surface staining against MHC II, permeabilization was omitted. Cells were blocked with 5% BSA (Sigma Aldrich, cat. no. 7030) for 1 h at room temperature. Primary antibodies (anti-OCT-1 (Invitrogen, cat. no. MA5-41265, 1:1000) and anti-MHC II (Cell Signaling, cat. no. 42594S, 1:200) were incubated for 1 h at room temperature. Anti-OCT-1 staining was furthermore incubated with 4 µg/mL anti-Rabbit Alexa Fluor 488 antibody (Invitrogen, cat. no. A-11008) for 1 h at room temperature. All samples were mounted with DAPI Fluoromount-G (SouthernBiotech, cat. no. 0100-20) and imaged with a Zeiss LSM Meta 710 confocal microscope. For MHC II staining, the pinhole was opened to allow for multi-layered imaging acquisition, as Z-stacking would have led to excessive signal decay. Images were analysed with ImageJ2 2.3.0/1.53f. For quantification of the nuclear OCT-1 signal, the nuclear area was defined manually according to the DAPI signal, followed by measuring the Alexa Fluor 488 signal. Integrated Fluorescence was extracted. Prism 9.2.0 was used for visualisation and statistical analysis.

### Flow Cytometry

For nuclei isolation, cells were incubated in NE1 buffer (20 mM HEPES-KOH pH 7.9, 10 mM KCl, 0.5 mM Spermidine, 0.1 % Triton X-100, 20 % glycerol, 1X cOmplete Proteinase Inhibitor) for 10 min on ice. Nuclei were fixed with 0.1 % formaldehyde (Thermo Scientific, cat. no. 28906) in PBS for 2 min at room temperature. Formaldehyde was quenched with 74 mM glycine. Nuclei were incubated with 1:1000 anti-OCT-1 antibody (Invitrogen, cat. no. MA5-41265) overnight at 4 °C. The next day, nuclei were incubated with 4 µg/mL anti-Rabbit Alexa Fluor 488 antibody (Invitrogen, cat. no. A-11008) for 1 h at room temperature. Nuclei were stained with 1 µg/mL DAPI (Sigma Aldrich, cat. no. D9542) in FACS buffer (1X PBS, 0.5 % FBS, 2 mM EDTA) to discriminate nuclei from debris in flow cytometry. Data was acquired on a BD FACSCanto II and analysed with FlowJo 10.7.2.

### Integration of our findings with patient data from TCGA

Lehmann et al. showed that TNBC can be divided into at least four subtypes based on patient data from TCGA^13^. Their provided scripts (https://github.com/TransBioInfoLab/TNBC_analysis) were used, modified and extended for our analysis. Data were retrieved from TCGA, and differential analysis was performed using limma. GSEA was performed using GSEA version 4.2.3 software on a pre-ranked gene list representing log2 fold change and FDR from the differential analysis. Default parameters were used for GSEA preranked, including gene set size between 15 and 500 and phenotype permutation at 1,000 times. As a gene set, the top 100 enriched genes in BMDM+TNBC vs BMDM were used and translated to human gene symbols. To investigate if Pou2f1 targets were enriched in one of the TNBC subtypes, we used the generated ranked lists from before and the Pou2f1 targets detected with our TOBIAS analysis as enriched in BMDM+TNBC translated to human symbols. Analysis was also performed using GSEA as above. As BL2 was enriched, we used only patient data with BL2 subtype and genes of interest detected in our study to generate a heatmap using ComplexHeatmap (ver. 2.14.0) and kmeans=4.

## Data analyses

### ChIP-seq data analysis

ChIP-seq data analysis was performed as previously described^10^. In short, Fastq files were trimmed from adapters using cutadapt^58^ (options: -q 20 -O 3, ver: 3.2) and aligned to a combined genome consisting of hg19 (Homo sapiens), dm6 (D. melanogaster) and mm10 (Mus musculus) genome assemblies with bwa-mem (ver. 0.7.17). BAM files were generated from the alignment with samtools view (options: - Sb -F2308 -q 10, ver: 1.10) and subsequently split by organisms to obtain three BAM files for each sample. Duplicated reads were marked and removed using Picard MarkDuplicates (ver: 2.25.7). For all organisms, the total sequencing coverage was quantified as the total number of unique reads aligning to the respective genome. Standard peak calling was performed for each sample using MACS2 (ver. 2.2.7.1) with default options. For each PDX model, mouse peak regions were considered positive if confirmed in two of four technical replicates (multi2) with bedtools (ver. 2.30.0) ‘multiIntersectBed’. All mouse-confirmed G4-ChIP–Seq peak files (multi2) of the nine models were merged (bedtools ‘merge’), and regions more than 99 bp long were retained to generate a single G4 DNA consensus file of 1578 G4 regions. Finally, the coverage of the samples was quantified using this consensus set (bedtools coverage). Reference normalisation factor estimation and mouse chromatin immunoprecipitation signal normalisation were performed as previously described^10^.

### CUT&Tag data analysis

CUT&Tag analysis was performed similarly to ChIP-seq analysis; however, the data was only aligned to mm10 as we used mouse BMDMs. Furthermore, peak calling was done using SEACR (https://github.com/FredHutch/SEACR).

### Differential binding analysis

Differential G4-binding analysis was employed to identify ΔG4Rs, as described using edgeR^10^ (R ver. 4.1.1, edgeR ver. 3.34.1). Initially, the library size and the *D. melanogaster*-normalised mouse read coverage within mouse G4 consensus regions were computed. Then, a generalised linear model with default parameters (negative binomial log-linear distribution of reading counts) was used to assess regions with differential binding signals. For CUT&Tag samples, the *D. melanogaster* normalization step was omitted. The differential binding analysis compared each PDX model to all the others. For each comparison, differential DNA G4 regions ΔG4R (that is, regions specifically present in a given PDX model) were defined as those satisfying the following criteria: log2(Cpm) ≥ 0.6 and false discovery rate < 0.05.

Similarity across all the 9 PDX ΔG4Rs was estimated using bedtools Jaccard. Jaccard indexes of all pairwise comparisons resulted in a nine-by-nine matrix. After the data was loaded into R, a pairwise intersection heatmap was generated with the following settings: plot type: corrplot; correlation coefficient: Spearman; order: hclust, hclust.method: ward.d2 (corrplot ver. 0.92).

### Genomic and G4-motif annotation and enrichment analysis of patient-derived tumour xenograft qG4-ChIP–seq peaks

PAVIS was used to annotate 1578 PDX G4 mouse consensus regions^59^. Fold enrichment analysis was performed as described^10^. The consensus peaks were ten times randomly shuffled across the genome. Fold enrichments were computed as the ratio between the fraction of overlaps with each genomic feature in the actual case versus the corresponding average random fractions.

### Differentially enriched G4 regions analysis in expressed gene promoters

Promoter transcription start site (TSS) coordinates, ±1 kb from the TSS, were generated for genes using the mm10 genome (GRCm38.102, ensemble.org); see section “Code Availability”. The fraction of ΔG4Rs overlapping high-, medium-, and low-expression gene promoters was estimated.

### RNA-Seq data analysis

For the PDX models, RNA-Seq data was obtained from EGA (ID: EGAD00001006307). For the BMDM samples, we generated RNA-seq data. After quality control using FastqQC (ver: 0.11.9), alignment was performed using STAR (ver:2.7.3a, options: twopassMode Basic, readFilesCommand gunzip -c, outSAMtype BAM SortedByCoordinate). For PDX models, a combined genome of hg19 and mm10 was used, whereas for BMDM samples, only mm10 was needed. Next, PDX models were split into mm10 and hg19 bam files using samtools. Then, htseq-count (ver: 2.0.1) and a mouse GTF file (GRCm38.102, ensemble.org) were employed to count the reads. After combining counts for all samples, differential analysis was performed in R. First, edgeR was used to compare each PDX model to all others. For BMDM, the control was compared to either BMDM+TNBC or BMDM+HR^+^. In the second step, DESeq2 (DESeq2 ver. 1.32.0) was also used for differential analysis. Heatmaps of the 30 most variable genes were generated after vst normalization using pheatmap (pheatmap ver. 1.0.12). GO terms were annotated using goana (part of limma package, limma ver. 3.48.3). Furthermore, GSEA was performed using GSEA version 4.2.3 software on a pre-ranked gene list representing log2 fold change and FDR of expression of BMDM+TNBC and BMDM+HR^+^. Default parameters were used for GSEApreranked, including gene set size between 15 and 500 and phenotype permutation at 1,000 times.

### Consensus^TME^

The tool consensus^TME 23^ (https://github.com/cansysbio/ConsensusTME) needs human symbols and counts per million. Therefore, a table from biomart (ensemble.org) was used to convert mouse IDs into human symbols, and the count matrix was converted into a DGEList object to apply Cpm. Next, the mean Cpm for each sample was calculated and used as input for consensusTME. The resulting normalised enrichment score was plotted as a heatmap.

### Transcription factor analysis using TOBIAS

TOBIAS (Transcription factor Occupancy prediction By Investigation of ATAC-seq Signal, https://github.com/loosolab/TOBIAS)^**25**^ can be used to identify TF footprints in ATAC-seq data. Here, it was used for the same purpose but with our G4-CUT&Tag data, as G4s predominantly form in open chromatin regions and are enriched in promoter regions of genes. We investigated TF footprints in the differentially enriched G4 regions of BMDM vs BMDM+TNBC or BMDM+TNBC vs BMDM+HR^+^ in their respective bam files. The analysis was performed as described for TOBIAS. TF footprints for all vertebrates were downloaded from (https://jaspar.genereg.net).

### IC Clusters

Murine differentially enriched genes of PDX models were associated with genes signifying integrative clusters (Ref 10 and Curtis et al. Nature 2012). Promoter coordinates of IC signature genes (adjusted P value < 0.05; log2(fold change) > 0.6) of each IC (1–10) were extracted. For each PDX model, the association of differentially enriched genes to IC signature promoters were quantified by computing the corresponding p-value (−log10(p-value)) from the Fisher’s exact test (intervene pairwise option fisher, intervene version 0.6.5). For highly significant associations, which resulted in p-values of 0, the −log10(p-value) was set to 300. -log10(p-values) were plotted as bars.

### Overlap of PDX model genes with *Pou2f1* binding sites enriched in BMDM+TNBC

*Pou2f1* binding sites enriched in ΔG4Rs of BMDM+TNBC were detected by TOBIAS. These regions +/-1 kb overlapped with region +/-2.5 kb of all genes detected in RNA-Seq of the PDX models in grouped by their expression into three groups (low, medium, and high expressed). To calculate fold enrichment, *Pou2f1* binding sites were randomly shuffled (10x) using bedtools shuffle and overlap with genes was calculated. For each PDX model, counts of overlaps were summed up per expression group and divided by the median gene length per expression group. The same procedure was applied to the shuffled data. To calculate fold enrichment, results from original *Pou2f1* bindings sites were divided by results from shuffled regions and plotted as a bar chart.

## Data availability

Raw and processed data files for BMDM experiments are available at the Gene Expression Omnibus (GEO, National Center for Biotechnology Information repository) and the accession number GSE218279.

The qG4-ChIP–seq data reported in this paper are available at GEO under accession number GSE152216. Gene-expression (RNA-seq) data of the PDTX models are available at the European Genome-phenome Archive under accession number EGAS00001001913.

## Code availability

Code is provided as supplementary data.

## Acknowledgements

We acknowledge the Cologne Center for Genomics for its support of NGS. We thank the Regional Computing Centre (RRZK) of the University of Cologne for its support. We recognise the imaging facility of the CECAD and the cell sorting facility of the CMMC for support. We would like to thank Oscar M. Rueda for his help in organising and obtaining the PDX RNA-seq data. The DFG CRC1399, DFG (HA 8562/4-1), CANTAR, the Center for Molecular Medicine Cologne and the Fritz-Thyssen Foundation support the Hänsel-Hertsch laboratory. Jan Herter and Grit S Herter-Sprie laboratories are supported by DFG (HE 6810/3-1 to JMH., HE 6897/2-1 to GSHS). We thank Dr Leo Kurian and his laboratory for providing access to their Bio-Rad CFX96 Touch Real-Time PCR Detection System.

## Contributions

R.H.-H. conceived this study and interpreted the results. R.H.-H. and P.H. wrote the manuscript. P.H. performed all experiments and interpreted the results. O.v.R. performed experiments. MNH analysed all data, interpreted results and created figures. MK prepared macrophages. GSHS, JMH, MK and MNH edited the manuscript.

---

**Supplementary Figure 1.**
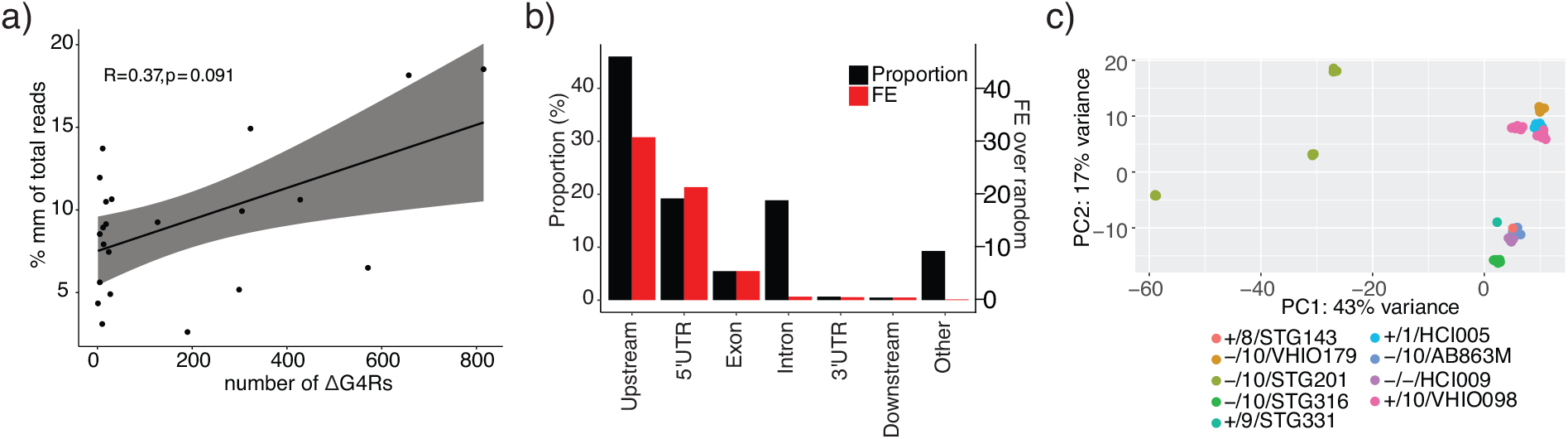
Supporting analyses for Figure 1. **a** Spearman correlation of the level of ΔG4Rs (x-axis) and percentage of mouse reads (y-axis) in PDX models. **b** Genome annotation of the nine PDX mm10 ΔG4Rs. Black bars: proportion of G4 regions in particular genomic annotation, red bars: fold enrichment over random genomic regions (n = 10 permutations). Data are presented as mean values. **c** PCA plot of RNA-seq data for the nine PDX models over random genomic regions (n = 10 permutations). Data are presented as mean values.

**Supplementary Figure 2.**
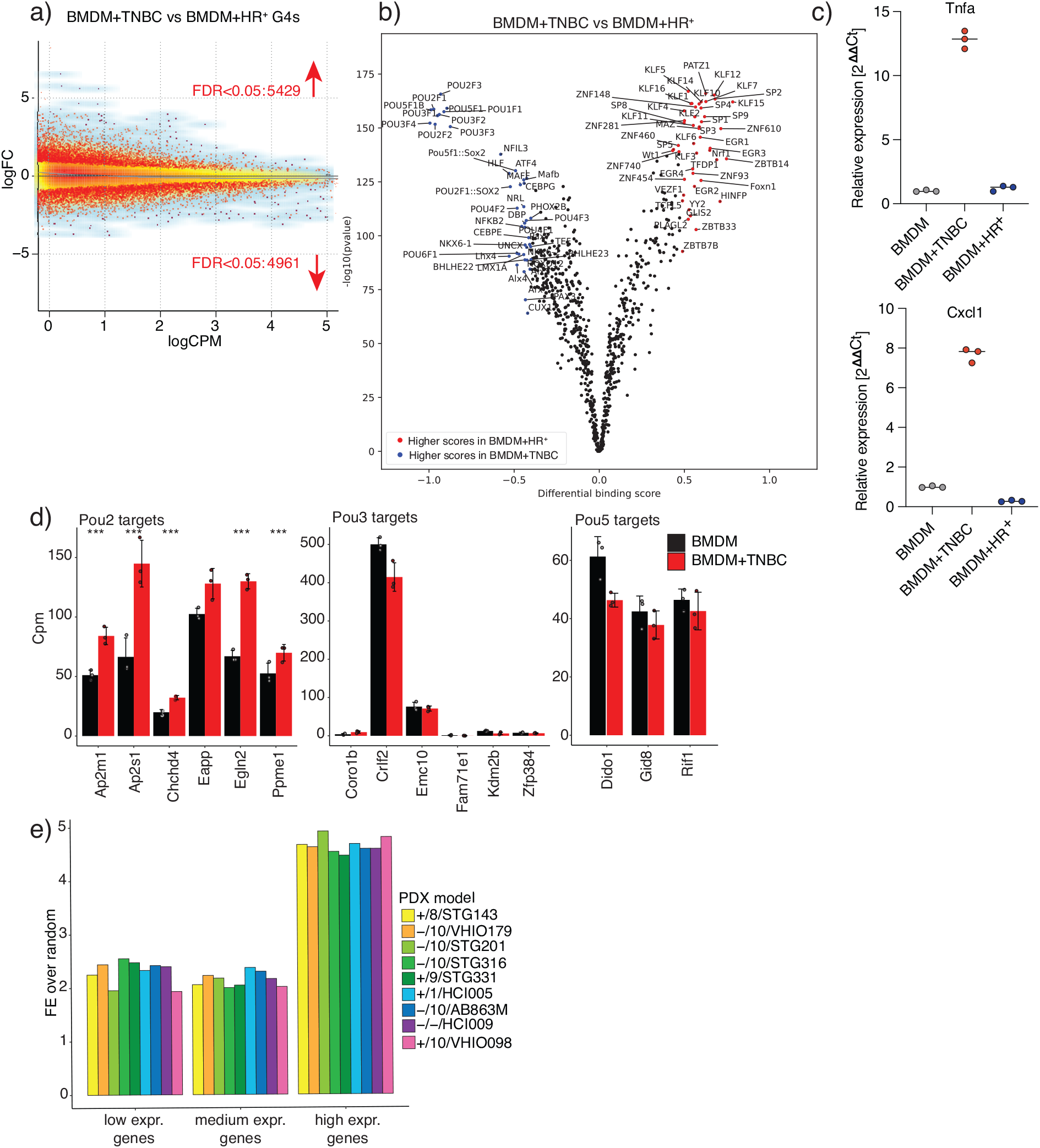
Supporting analyses for Figure 2. **a** Bland–Altman plot (MA) of BMDM cultured in TNBC-conditioned medium vs BMDM cultured in HR^+^-conditioned medium. Red dots= differentially enriched. Blue= smooth scatter, grey line= Smooth Curve Fitted by Loess. **b** Same as Fig. 2d but for BMDM+TNBC vs BMDM+HR^+^. **c** qPCR for *Tnfa* and *Cxcl1*. **d** mean vst normalised expression and standard deviation of different *Pou* family member targets in RNA-seq data of BMDM and BMDM+TNBC. *** p-value < 0.001 from the differential analysis. **e** Mean fold enrichment over random of overlaps of PDX genes grouped by expression with *Pou2f1* binding sites enriched in ΔG4Rs of BMDM+TNBC.

**Supplementary Figure 3.**
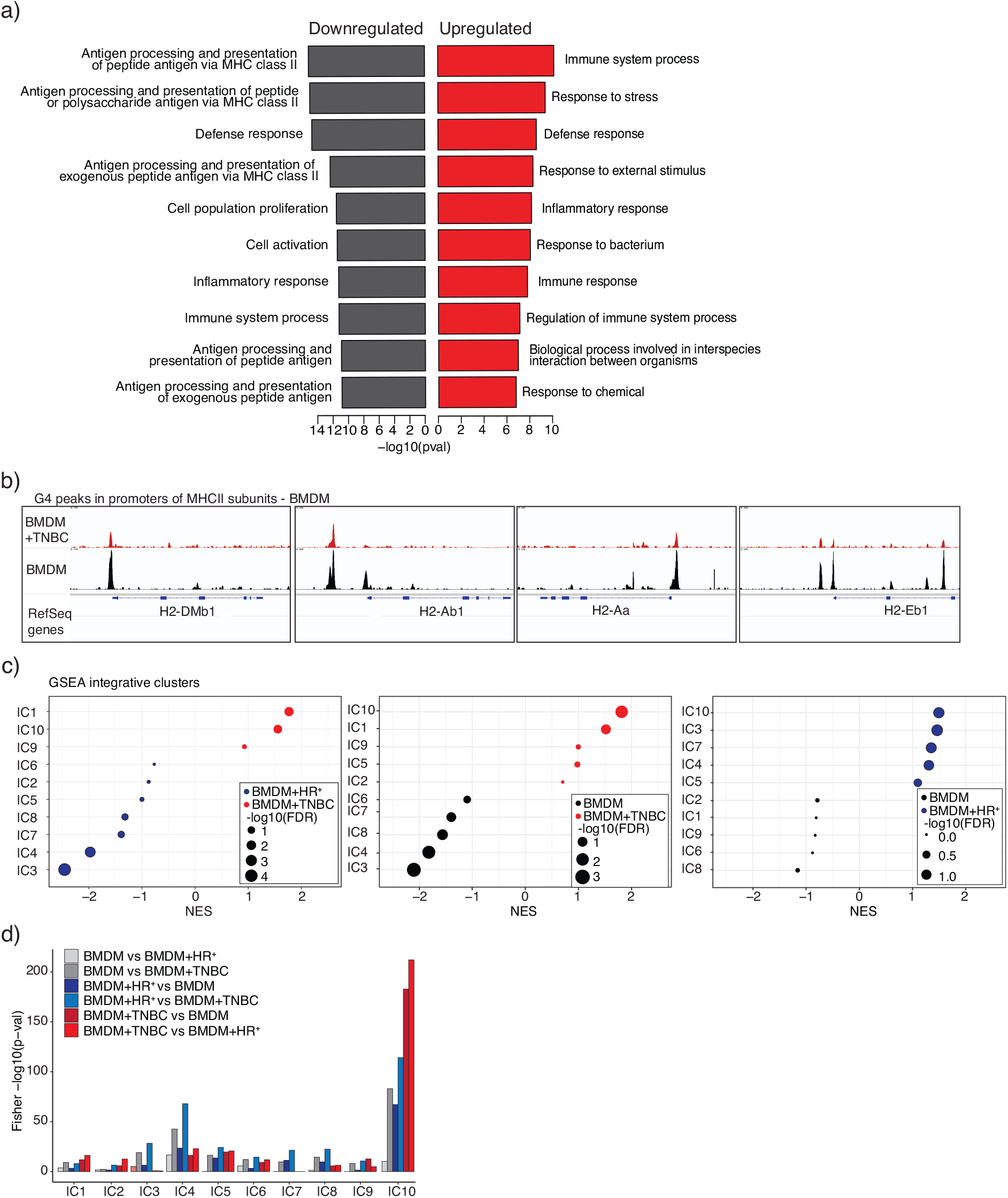
Supporting analyses for Figure 3. **a** Gene ontology (GO) term analysis of differentially enriched genes for BMDM and BMDM+TNBC conditions. The y-axis depicts significance, and the x-axis the top enriched GO terms. **b** IGV snapshots of BMDM (black) and BMDM+TNBC (red) G4 signal in MHC class II subunits. **c** GSEA preranked of BMDM+TNBC vs BMDM+HR^+^, BMDM vs BMDM+TNBC or BMDM+HR^+^, respectively, and ICs as gene sets. **d** as in Fig. 3e: - log10(p-value) of Fisher’s exact test extracted from overlaps of the integrative cluster (IC) signature genes (molecular subtypes of breast cancer) with differentially enriched genes revealed in all combinations of BMDM vs BMDM+TNBC vs BMDM+HR^+^, respectively.

